# Neuropathic injury drives a generalized negative affective state in mice

**DOI:** 10.1101/2022.11.10.515959

**Authors:** Makenzie R. Norris, John Bilbily, Léa J. Becker, Gustavo Borges, Yu-Hsuan Chang, Samantha S. Dunn, Manish K. Madasu, Ream Al-Hasani, Meaghan C. Creed, Jordan G. McCall

## Abstract

Neuropathic pain causes both sensory and emotional maladaptation. Preclinical animal studies of neuropathic pain-induced negative affect could result in novel insights into the mechanisms of chronic pain. Modeling pain-induced negative affect, however, is variable across research groups and conditions. The same injury may or may not produce robust negative affective behavioral responses across different species, strains, and laboratories. Here we sought to identify negative affective consequences of the spared nerve injury model on C57BL/6J male and female mice. We found no significant effect of spared nerve injury across a variety of approach-avoidance, hedonic choice, and coping strategy assays. We hypothesized these inconsistencies may stem in part from the short test duration of these assays. To test this hypothesis, we used the homecage-based Feeding Experimentation Device version 3 to conduct 12-hour, overnight progressive ratio testing to determine whether mice with chronic spared nerve injury had decreased motivation to earn palatable food rewards. Our data demonstrate that despite equivalent task learning, spared nerve injury mice are less motivated to work for a sugar pellet than sham controls. Further, when we normalized behavioral responses across all the behavioral assays we tested, we found that a combined normalized behavioral score is predictive of injury-state and significantly correlates with mechanical thresholds. Together these results suggest that homecage-based operant behaviors provide a useful platform for modeling nerve injury-induced negative affect and that valuable pain-related information can arise from agglomerative data analyses across behavioral assays - even when individual inferential statistics do not demonstrate significant mean differences.

## Introduction

Chronic pain is a debilitating disease, severely impacting patients in their activities of daily living. In addition to sensory impairments, about half of chronic pain patients present with affective disorders, such as depression or anxiety[4,68,81]. These comorbidities exacerbate the burden of chronic pain and complicate the therapeutic management of each condition[5,31]. Therefore, to advance novel therapeutic strategies it is urgent to gain better insights into the pathophysiology of pain-induced negative affect. To that end preclinical models can be powerful tools.

The comorbidity between chronic pain and affective disorders can, to some extent, be modelled in rodents[2,15,17,23,44,49]. Thus, alterations in coping strategies, approach-avoidance, or hedonic behaviors[26,39,52,53,62,63,74,86], as well as decreased motivation, learning, and cognition[6,21,53,56,57,60,69,72] have been reported in preclinical studies. However, pain-induced negative affect is sometimes difficult to replicate across labs, and several groups consistently reported the absence of association between chronic pain and affective disorders[37,65,66,82]. Potential explanations for these discrepancies are diverse. Variability likely arises from the pain model used, those relying on nerve ligation or spinal cord injury being more variable [44,55]. Animal subject sex, experimenter sex, and the particular rodent strain used, are all known to separately affect pain and emotional processing[1,34,44,70,77,82], but likely also impact pain-induced negative affect. Importantly, the time-dependent nature of the chronic pain-induced affective consequences is a critical variable[2,84,85], and the absence of chronic pain-induced emotional deficits often occur in early-stage behavioral testing[44]. Furthermore, the acute nature of the behavioral tests assessing affective states in rodents, usually lasting less than a few minutes, should be considered. Affective states fluctuate over time[11], therefore using short duration behavioral assays may render emotional deficits challenging to observe. Surprisingly, this question has not been extensively tested in the preclinical pain-induced negative affect field.

Here, we hypothesized that longer duration assays and later testing time points might be more suitable to highlight the negative affective consequences of neuropathic pain in C57BL/6J mice. To test this hypothesis, we tested mice 7 to 26 weeks following spared nerve injury (SNI) in an extensive battery of affective behavioral assays. While we observed robust, chronic SNI-induced mechanical hypersensitivity, we found no effect on multiple, brief approach-avoidance, hedonic value, and coping strategy assays. To determine if longer access assays could be more appropriate, we used the homecage-based Feeding Experimentation Device version 3 (FED3)[58] to conduct multiple operant tasks. We tested mice in short access (1 hour) fixed ratio-1 (FR1) and fixed-ratio-3 (FR3) paradigms as well as a 12-hour, overnight progressive ratio schedule. Our results indicate that SNI decreases the motivation of mice to retrieve palatable food, suggesting that progressive ratio schedules may be a critical preclinical behavioral tool for better understanding of pain-induced negative affect. Finally, we normalized behavioral scores across assays[35] to more broadly understand the impact of SNI on the negative affective behaviors. Importantly, these results showed that performance on tests without statistical significance still strongly correlate with mechanical hypersensitivity and can provide a single metric for assessing the injury state.

## Methods

### Animals

Adult male and female C57BL/6J mice were used from age 8-34 weeks. Mice in spared nerve injury experiments were originally sourced from The Jackson Laboratory (Bar Harbor, ME, USA) and bred in-house in a barrier facility in another building. These animals were transferred to a holding facility adjacent to the behavioral space between 4-6 weeks of age. Mice were then left undisturbed except for necessary husbandry to habituate to the new facility until 8 weeks of age. All mice were group-housed, given *ad libitum* access to standard laboratory chow (PicoLab Rodent Diet 20, LabDiet, St. Louis, MO, USA) and water, and maintained on a 12:12-hour light/dark cycle (lights on at 7:00 AM). All experiments and procedures were approved by the Institutional Animal Care and Use Committee of Washington University School of Medicine in accordance with National Institutes of Health guidelines.

### Blinding and Randomization

Complete blinding with SNI mice is not possible because the injury causes noticeable postural changes that makes these animals visually distinct from sham controls. However, animals were blindly tested prior to SNI surgery and randomized to SNI or sham condition by cage. All video analyses were conducted automatically in Ethovision without human intervention.

### Spared Nerve Injury (SNI)

The surgical procedure for the SNI-induced model of neuropathic pain was performed as described previously[20]. Mice were anesthetized with 3% isoflurane and right hind limb shaved and disinfected with 75% ethanol and betadine. A 10-15 mm incision was made in the skin proximal to the knee to expose the biceps femoris muscle. Separation of the muscle allowed visualization of the sciatic nerve trifurcation. The common peroneal and tibial branches were ligated with 6-0 silk suture (Ethicon Inc., Raritan, NJ, USA) and 1 mm of nerve was excised distal to the ligature, leaving the sural branch intact. Following wound closure mice were allowed to recover on a table warmed to 43°C for 10-20 minutes prior to being returned to their home cage. Sham surgeries were identical to the SNI procedure without the ligation, excision, and severing of the peroneal and tibial branches of the sciatic nerve. Wound clips were removed from the healed incision after the first testing was completed on post-operative day 7.

### Mechanical sensitivity (von Frey)

Mechanical withdrawal thresholds were determined using von Frey filaments (Bioseb, Pinellas Park, FL,USA). Mice were acclimated for 2 hours on an elevated wire mesh grid in 5-inch diameter plexiglass cylinders wrapped in black opaque plastic sheets. Von Frey filaments were then applied to the lateral aspect of the hind paw[27], at a force ranging from 0.02 g to 3.5 g using the up-down method as described previously[18]. Each von Frey filament stimulation for each mouse was separated by 2 minutes. 50% withdrawal threshold was calculated as previously described[18]. All animals were tested before SNI surgery to determine baseline mechanical withdrawal thresholds and then once per week for the next six weeks and then again at week 26 following the completion of the final behavioral tests.

### Open field test (OFT)

The open field test was performed as previously described [59]. Mice were habituated to the test room environment for two hours prior to open field testing. The OFT was performed in a 50×50 cm enclosure. The center was defined as a square of 50% of the total OFT area. Mice were left to explore the apparatus for 20 minutes and the time spent in center was determined using Ethovision behavioral tracking software (Noldus). Overhead lighting during habituation and open field testing were stabilized between 7-9 lux as measured by a luxmeter.

### Elevated plus maze (EPM)

Mice were habituated to the test room environment for two hours prior to elevated plus maze testing. The test was performed on a +-shaped apparatus elevated 50 cm above the floor with two oppositely positioned closed arms, two oppositely positioned open arms, and a center area. The mice were placed in the center area and allowed to freely explore for 15 minutes. The center point of the mouse was used for behavioral tracking. Time spent in open arms was determined over 15 minutes using Ethovision behavioral tracking software (Noldus). Lighting during habituation and elevated plus maze testing were stabilized between 7-9 lux.

### Sucrose preference

The sucrose preference testing protocol was adapted from Liu et al. 2018[51]. First, mice were introduced to the two-bottle setup (1% w/v sucrose and water) in their home cage for 48 hours, this is defined as acclimation. 24 hours into acclimatation the bottles were switched to avoid inducing a side-bias. Next, to adapt the mice to the testing setup, all animals were single-housed, moved to the experimental room and given access to two-bottle setup on the cage top for 24 hours, this is defined as adaptation. After 24 hours of adaptation, mice were returned to their home cage, taken back to their holding room and water deprived for three hours. Finally, mice were returned to the test setup (moved to experimental room and singly housed with the two-bottle setup) two hours after the start of the dark cycle and tested for 12 hours. Two baseline test sessions were conducted before the experimental testing. All testing sessions occurred after three hours of water deprivation. Sucrose preference scores were calculated as the amount of sucrose intake/total liquid intake*100 during the experimental test. Baseline test data was not quantified. Mice were given *ad libitum* access to chow during the whole testing period.

### Tail Suspension Test (TST)

After 30-minute habituation to the testing room, mice were suspended 35 cm above an experimental bench by their tail for 6 minutes with a clear plastic cylinder placed at the tail base to prevent tail climbing behavior. Videos of tail suspension were taken using a Google Pixel 3 XL (Google, Mountain View, CA, USA) smartphone. Only the last 5 minutes of the 6-minute videos were considered when scoring for immobility behavior as an indication of learned helplessness. Passive swaying was not included in immobility time.

### Chocolate preference

Mice were given 12-hour access to chow and chocolate pellets in their home cage during the dark cycle for three nights. This habituation consisted of placing one gram of each pellet type at opposite ends of the cage (Test Diet catalog #1811251 & #1811256). On the test day, home cages with bedding removed were used as test enclosures. Two plastic weigh boats were taped to opposite ends of the empty home cage. Each weight boat was filled with one gram of either chocolate or chow pellets. Mice were then individually placed in the test enclosure for three hours. After the three-hour test, mice were placed back in their home cage and weigh boats containing either chocolate or chow pellets were weighed. Percent chocolate preference was defined as grams of chocolate pellet eaten/ (total grams of chocolate and chow pellet eaten) multiplied by 100.

### Nest building

Nest building was assessed as previously described [25]. Briefly, at the onset of the dark cycle, mice were singly housed in home cages with 3 g of cotton-based nestlet and access to *ad libitum* food and water. 16 hours later, mice were placed back into the original group housed home cages and nests were scored as previously described[25]. Intact nestlets were weighed. Mice received scores between 1 and 5. Factors considered in nestlet scores include: weight of remaining intact nestlets and overall shape of nests. Specific parameters contributing to overall nest score were followed as clearly outlined by Deacon, 2006[25]. Nest scores were blindly assessed using two scorers. Each investigator scored nests independently and compared nest scores to each other. Discrepancies between scores were discussed and final scores were agreed upon between both scorers.

### Social interaction

Home cage social interaction was performed as previously described [36,67]. A test mouse was placed into a new home cage with the lid removed and allowed to habituate to the novel environment for ten minutes. After ten minutes of habituation, an age- and sex-matched C57BL/6J stranger mouse was placed into the same home cage as the test mouse for three minutes, this was considered the test session. After three minutes of testing, both mice were returned to their original home cages. Interactions were scored for the first 2 minutes of the testing session. Videos of the three-minute testing session were acquired using a Google Pixel 3 XL(Google, Mountain View, CA, USA) smartphone and scored manually. Examples of behavior scored as interactions were active attention towards stranger mouse, walking towards stranger mouse, touching, and licking. Converse investigation of test mouse by stranger mouse was not included in the overall interaction time values.

### Two chamber social preference

The three chamber social preference[75] test was adapted to two chambers as previously described[76]. Test mice were placed in the middle of opaque black boxes measuring 54 cm length x 26 cm width x 26 cm height split into two equal chambers with two metal mesh enclosures centered in each chamber. Mice were allowed to freely explore the test setup without the presence of a stranger mouse for 10 minutes immediately prior to experimental testing, this was considered a baseline test. Following baseline testing, mice were removed, and a sex- and age-matched stranger mouse was placed into one of the metal mesh enclosures centered in one side of the black box. Mice were then allowed to freely explore the test apparatus with the presence of a stranger mouse on one side for 10 minutes. Interaction zones were defined as double the diameter surrounding wire enclosures. Automatic behavioral tracking was done using Ethovision.

### Operant behavior using Feeding Experimentation Devices

FED3s[58] were equipped with two nosepoke holes (active and inactive). Active nosepoke holes were reinforced with an accompanying 4 kHz, 300 ms tone and activation of a light bar directly underneath pellet receptacle. All animals were singly housed in fresh homecages with a cotton isopads purchased from BrainTree Scientific Inc. (Braintree, MA). All testing occurred consecutively overnight beginning at the start of the dark cycle for approximately 15 hours, to standardize across mice, only the first 12 hours were analyzed. Fixed-ratio 1 (FR1) testing was conducted until mice had 75% correct nosepokes for 3/5 testing sessions. Once mice met this criterion for FR1 testing, they advanced to three consecutive overnight sessions of fixed-ratio 3 (FR3) testing. Upon completion of FR3 testing mice went through a single overnight progressive ratio (PR) test on an exponential schedule. Chocolate sucrose pellets used for all FED3 testing sessions were purchased from LabDiet (St. Louis, MO, catalog number 1811149 (5TUT)). “Meal size” is a measure intended to estimate feeding patterns where multiple pellets are received from the FED3 in quick succession. Meal size was determined by setting a maximum interpellet interval of five minutes with a minimum meal size of one pellet.

### *Emotionality z-score* calculation

To capture the effect of SNI on mice emotional reactivity we computed emotionality z-scores as previously described [35]. First, individual z-score values were calculated for each test parameter using the following formula:

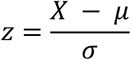

where X represents the individual data for the observed parameter while μ and σ represent the mean and standard deviation of the control group. The directionality of scores was adjusted so that increased score values reflect increased dimensionality (locomotion, avoidance, coping, nesting or social behaviors and motivation). For instance, decreased time spent in center in the OFT or decreased time spent in the open arms of the EPM, were converted into positive standard deviation changes compared to group means indicating increased avoidance behaviors. Conversely, increased immobility time in the TST was not converted since it is a direct indicator of increased helplessness behaviors. Z-score for each mouse in each test were then calculated by averaging the z-score obtained for each parameter. For instance, Z_OFT_ was computed as follow:

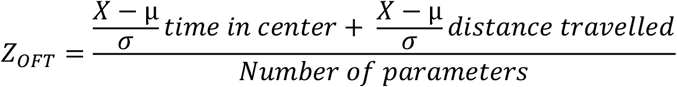

The same was applied to the other tests using the following behavioral parameters: OFT: time in center and distance travelled; EPM: time in open arm and distance travelled; Sucrose preference test: percentage of sucrose consumption; TST: immobility time; PR: breakpoint and the number of leftover pokes per animal after its breakpoint; FR: pellets earned in each of the 3 test sessions; SI: interaction frequency and interaction time; Nest test: percentage of nest torn and nest score; Two-chamber social preference test: time spent in stranger side; Chocolate preference: percentage of chocolate consumption.

Finally, z values obtained for each test were averaged to obtain a single emotionality score for each cohort:

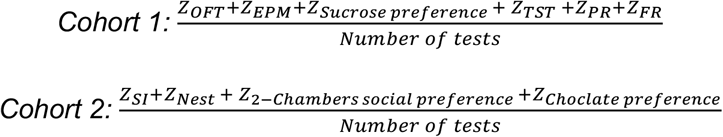

### Statistics and data analysis

All data are expressed as mean ± SEM. In data that were normally distributed, differences between groups were determined using independent t-tests or one-way ANOVA, or two-way ANOVAs followed by post hoc comparisons if the main effect was significant at p < 0.05. In cases where data failed the D’Agostino and Pearson omnibus normality test or was ordinal by nature, non-parametric analyses were used. Statistical analyses were conducted using Prism 9.0 (GraphPad). CSV files generated by FED3 were processed and plotted using FED3VIZ graphical user interface to generate plots. FED3VIZ is a custom open-source graphical program for analyzing FED3 data[58]. FED3VIZ provides plotting and data output for visualizing different aspects of FED3 data, including pellet retrieval, poke accuracy, pellet retrieval time, delay between consecutive pellet earnings (inter-pellet intervals), meal size, and progressive ratio breakpoint. Correlation data was assumed to follow a Gaussian distribution and computed using a Pearson correlation coefficient.

## Results

### Spared nerve injury induces long-term ipsilateral mechanical hypersensitivity and delayed onset contralateral mechanical hypersensitivity

The spared nerve injury (SNI) model of neuropathic pain has been shown to reliably induce ipsilateral mechanical hypersensitivity across many studies[10,71]. Here we performed SNI or sham surgeries and used the von Frey assay to assess mechanical withdrawal thresholds (**Figure 1A&B**). As expected, SNI induced hindpaw mechanical hypersensitivity ipsilateral to the injured nerve. Similar assessment of the contralateral side showed a gradual decrease in mechanical withdrawal threshold in SNI mice during week four that returned to baseline by week six. When tested again during week twenty-six we saw a profound allodynic effect in the contralateral hindpaw of SNI mice like the hypersensitivity seen on the ipsilateral side (**Figure 1C&D**). In summary, the spared nerve injury model induced robust prolonged mechanical hypersensitivity in the ipsilateral paw. Interestingly, our results show some evidence of contralateral sensitization as a result of a chronic injury state indicated by significant mechanical hypersensitivity of the noninjured limb 26 weeks after surgery. Following six weeks SNI-induced mechanical hypersensitivity, we next tested two cohorts of animals across a battery of tests to determine the extent to which SNI changes affective behaviors in C57BL/6J mice (**Table 1**).

**Figure 1:**
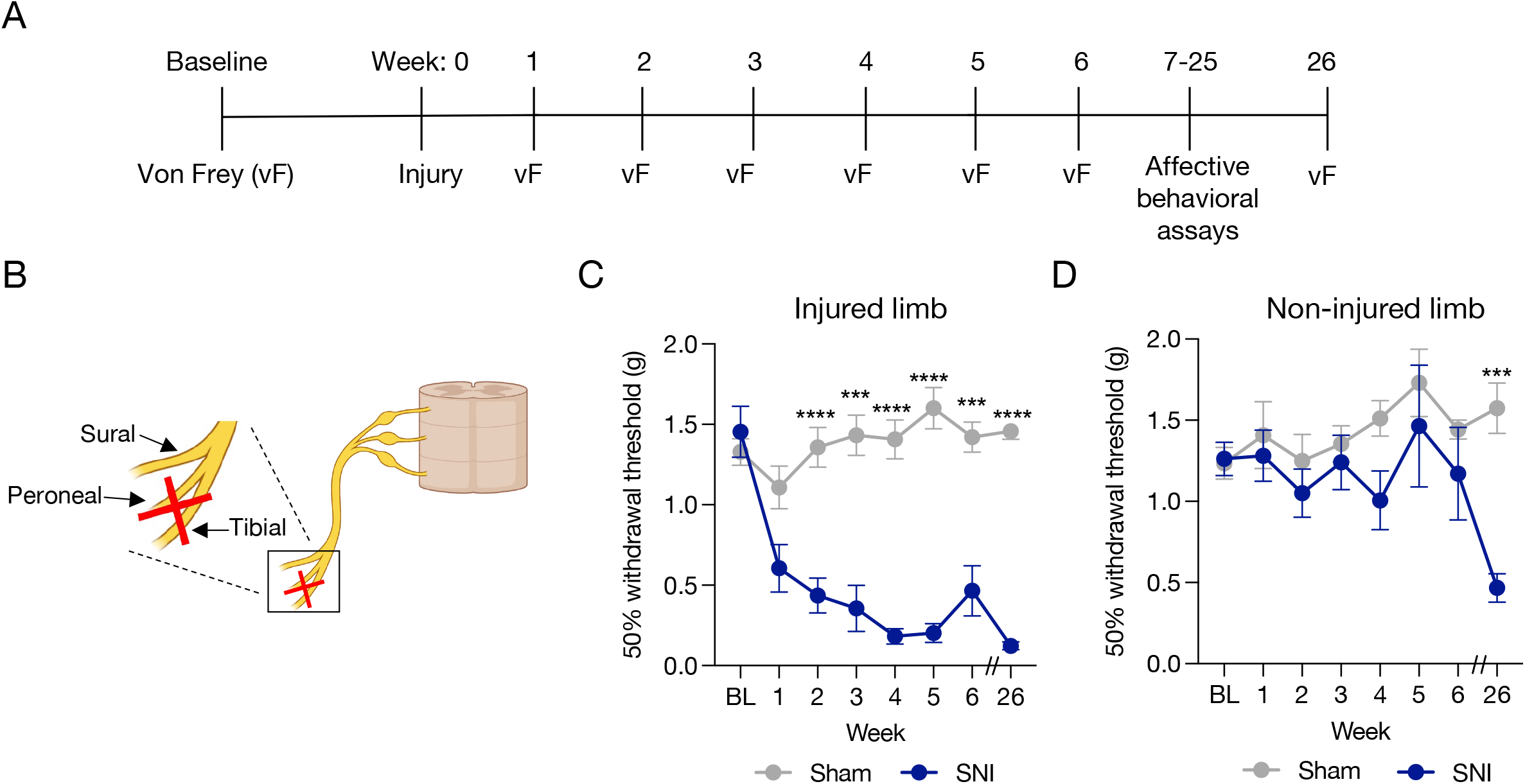
Spared nerve injury induces long-term mechanical hypersensitivity. (**A**) Experimental timeline. (**B**) Cartoon of spared nerve injury procedure. (**C**) SNI significantly decreases ipsilateral 50% mechanical withdrawal threshold through week 26 (Two-way Repeated Measures ANOVA, Sidak’s post hoc; Injured limb Sham vs. SNI F(1,41)=351.0). (**D**) SNI produces some contralateral hypersensitivity, particularly after prolonged injury (Two-way ANOVA, Sidak’s post hoc: Non-injured limb Sham vs. SNI F(1,41)=14.25). Data represented as mean +/− SEM; n=18-20/group; *p<0.05 ***p<0.001 ****p<0.0001.

**Table 1.** Table identifies specific behavioral assays assigned to cohorts 1 and 2 of mice. Abbreviations: FR1, fixed-ratio 1; FR3, fixed-ratio 3; PR, progressive ratio.

| Cohort | Construct assessed | Behavior test | Primary outputs |
| --- | --- | --- | --- |
| 1 | Mechanical sensitivity | Von Frey | 50% withdrawal threshold |
| 1 | Approach-avoidance | Open Field | % time in center<br>distance travelled |
| 1 | Approach-avoidance | Elevated plus maze | % time in open arms<br>distance travelled |
| 1 | Hedonic value | Sucrose preference | % sucrose preference |
| 1 | Learned helplessness | Tail Suspension | time spent immobile |
| 1 | Learning & Motivation | Operant tasks<br>FR1<br>FR3<br>PR | All:<br>pellets earned<br>% correct<br>retrieval time<br>meal size<br>FR1:<br>days to criteria<br>FR3:<br>time to 35 pellets<br>interpellet interval<br>PR:<br>breakpoint<br>pokes leftover<br>interpellet interval |
| 2 | Hedonic value | Chocolate preference | % chocolate preference |
| 2 | Innate nest building | Nestlet test | % nestlet torn<br>nest score |
| 2 | Socialization | Social interaction | interaction frequency<br>total interaction time |
| 2 | Socialization | Social preference test | time spent in stranger side |

### Chronic neuropathic injury does not alter approach-avoidance behaviors

Mice tend to avoid open areas, prefer darker spaces, and stay closer to walls (i.e., thigmotaxis). These natural behaviors present mice with an approach-avoidance challenge upon exposure to open areas. Quantifying this conflict is a common method used to assess behaviors thought to be relevant to anxiety-like behaviors in mice[19,50,59]. Here we quantify approach-avoidance conflict using the open field test (OFT) and elevated plus maze (EPM). We found that six weeks of SNI injury did not alter time spent in the center or distanced travelled in the open field (**Figure 2A-C**). Likewise, six weeks of SNI did not alter time spent in open arms of the elevated plus maze (**Figure 2D&E**), nor distance traveled in the whole maze (**Figure 2F**). This suggests six weeks of chronic neuropathic injury does not significantly alter the conflict inherent in approach-avoidance behaviors or gross locomotor activity. The lack of changes to overall locomotion in both assays suggests SNI does not induce ambulatory effects that could specifically confound these interpretations.

**Figure 2:**
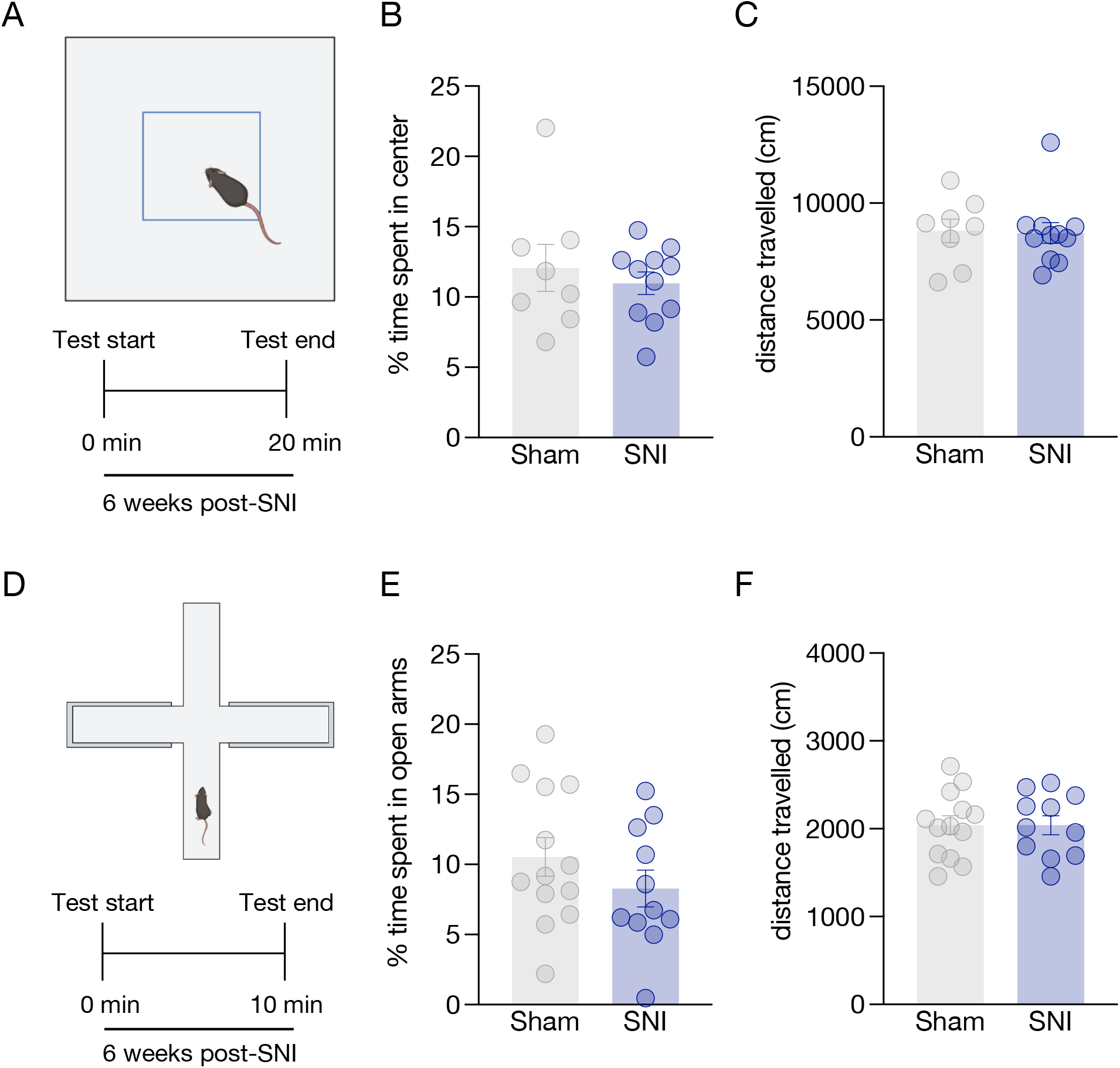
Chronic spared nerve injury does not alter approach-avoidance behaviors. (**A**) Cartoon of open field test behavioral apparatus and experimental timeline. SNI did not alter % time spent in center zone (**B**) or total distance travelled (**C**). Data represented as mean +/− SEM; n=8-11/group; unpaired t-test (Sham vs. SNI) (B) t=0.644; p=0.5279, (C) t=0.5663; p=0.5786, (D) t=0.1358; p=0.8935. (**D**) Cartoon of elevated plus maze behavioral apparatus and experimental timeline. SNI did not alter % time spent in open arms (**E**) or total distance travelled (**F**). Data represented as mean +/− SEM; n=11-13/group; unpaired t-test (Sham vs. SNI) (F) t=1.180; p=0.2533, (G) t=1.119; p=0.2751, (H) t=0.9239; p=0.3685.

### Chronic spared nerve injury did not alter hedonic value or suspension-induced learned helplessness

To assess depressive-like behavioral phenotypes during chronic neuropathic injury we used the sucrose preference test, tail suspension test, and a palatable food choice assay. Preference for rewards is a conserved phenomenon among mammalian species. In humans, depression is commonly associated with a loss of hedonic value or a lack of positive response to normally rewarding stimuli[13]. In rodents we can model this phenomenon by presenting mice with a choice between water and sucrose solution and measuring the animal’s preference for either solution. We found that eight weeks of spared nerve injury did not alter preference for a sucrose liquid solution (**Figure 3A&B**). In a separate cohort of animals, we tested a preference for chocolate pellets (**Figure 3E)** over normal chow. As in the sucrose preference test, eleven weeks of SNI also did not alter the choice of palatable foods (**Figure 3F)**. Together these results suggest longterm spared nerve injury does not alter innate hedonic value. The tail suspension test acts as a measure of learned helplessness and is commonly used as a screening method for potential antidepressant therapeutics[22]. In the tail suspension test, nine weeks of SNI injury had no effect on immobility time suggesting chronic spared nerve injury does not alter mobility during suspension in mice (**Figure 3C&D**). Together these results suggest spared nerve injured mice do not develop either anhedonia, as indicated by the sucrose preference and palatable food preference tests, or learned helplessness phenotypes, as measured with immobility time in the tail suspension test.

**Figure 3:**
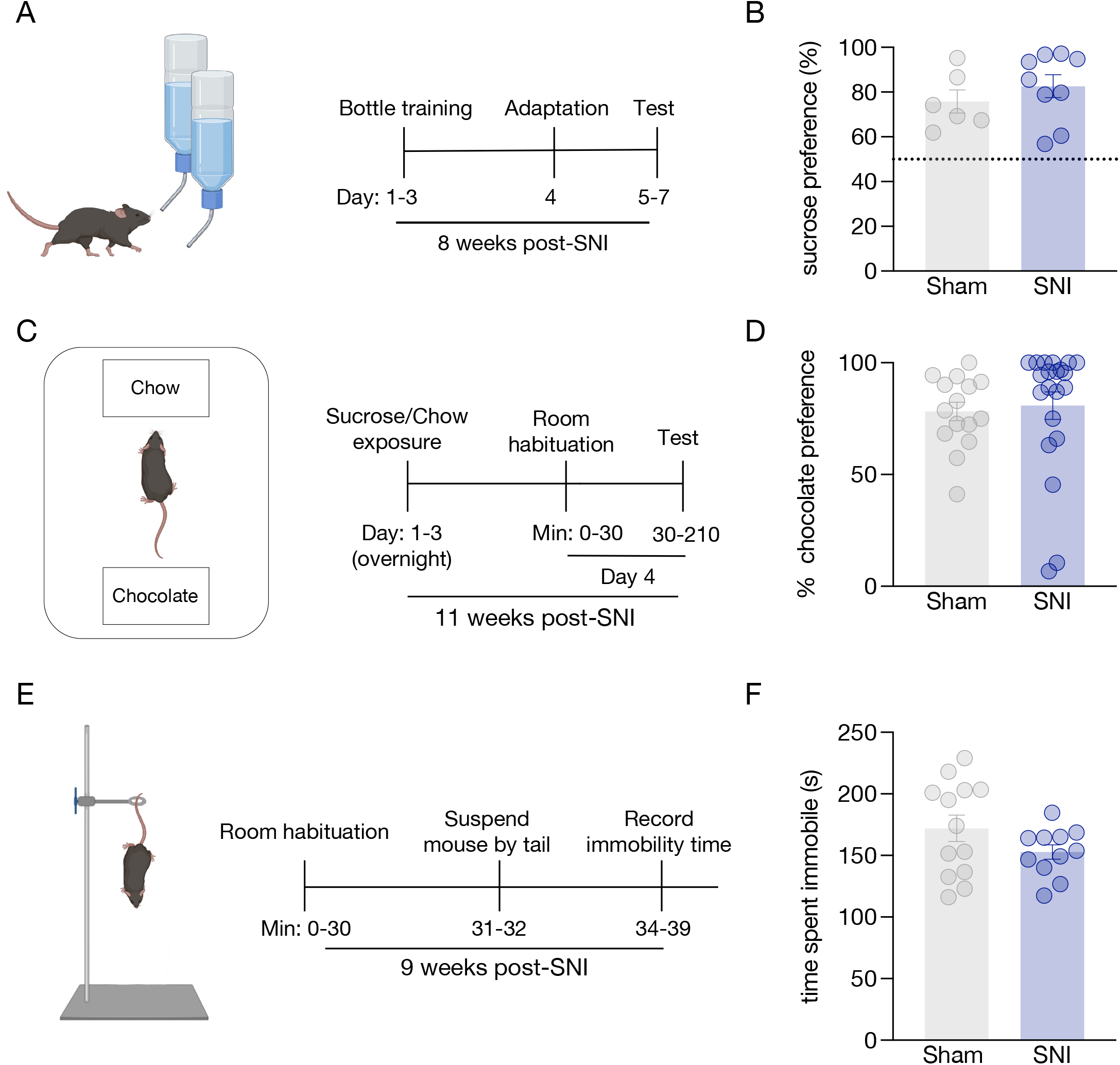
Chronic spared nerve injury does not alter hedonic choice or behavior during tail suspension. (**A**) Cartoon of sucrose preference test and experimental timeline. (**B**) SNI did not alter preference of sucrose relative to water. Data represented as mean +/− SEM; n=6-9/group; unpaired t-test (Sham vs. SNI) t=1.180; p=0.2533. (**C**) Cartoon of food preference test and experimental timeline. (**D**) SNI did not alter preference of chocolate pellets relative to chow. Data represented as mean +/− SEM; n=15-21/group; unpaired t-test (Sham vs. SNI) t=0.3255; p=0.7468. (**E**) Cartoon of tail suspension test and experimental timeline. (**F**) SNI did not alter time spent immobile in seconds. Data represented as mean +/− SEM; n=11-13/group; unpaired t-test (Sham vs. SNI) t=1.509; p=0.1391.

### Chronic spared nerve injury did not disrupt nest building behavior

Innate nesting behavior has been used as a proxy of general well-being in mice [25,33] and occasionally suggested as a model of obsessive-compulsive disorder-related behaviors[54]. Briefly, higher quality nests tend to be associated with mouse health and variability between nests built may provide insights into differences into whether this daily, cognitive task remains intact. (**Figure 4A&B)**. Here we show that 14 weeks of SNI injury does not influence the % of total nestlet torn (**Figure 4C**) or alter overall composite nestlet scores (**Figure 4D**). These data suggest 14 weeks of SNI does not disrupt a proxy of mouse occupational health as indicated by innate nest building behavior.

**Figure 4:**
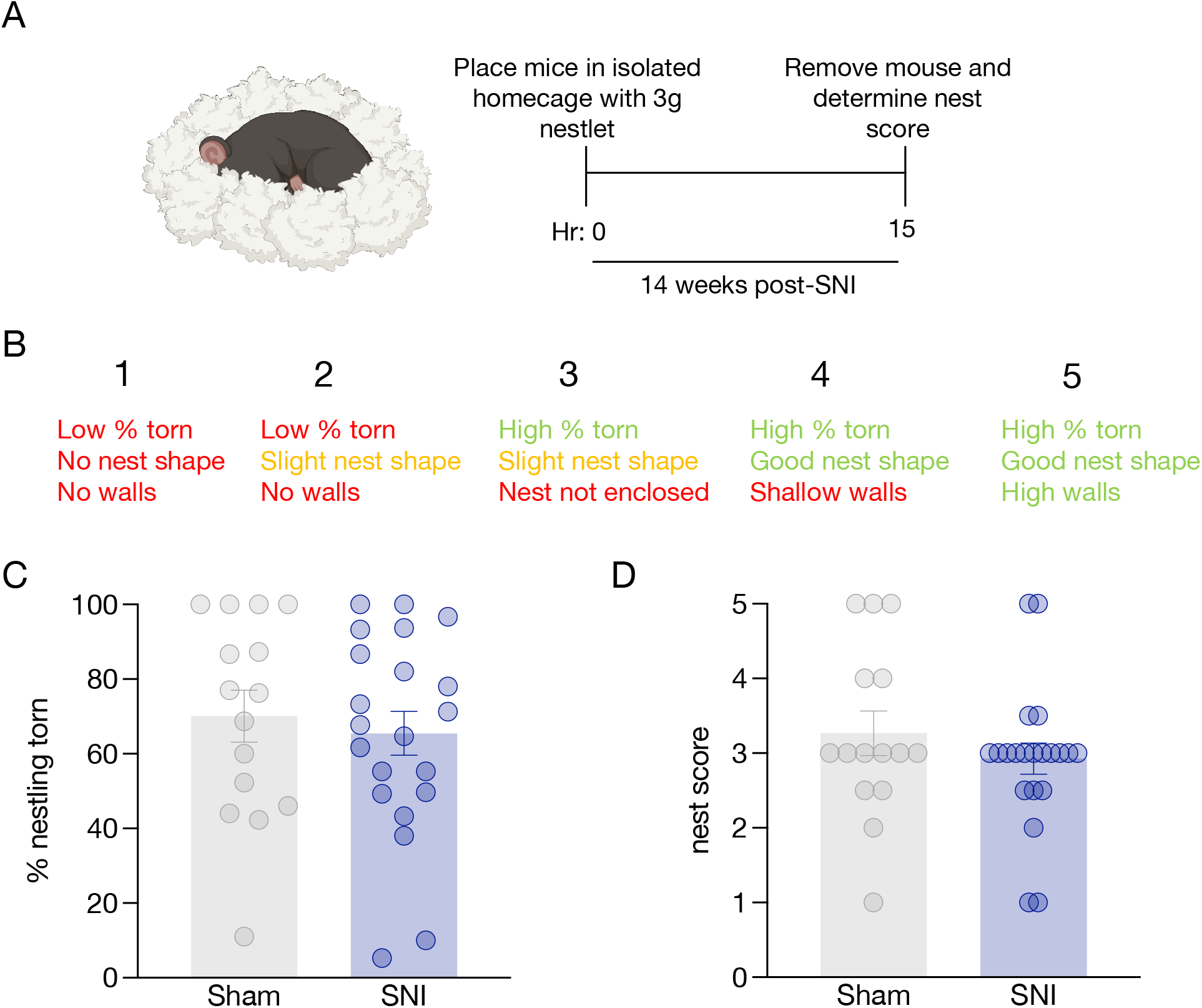
Chronic spared nerve injury does not disrupt innate nest building behavior. (**A**) Experimental timeline. (**B**) Nest building scores and associated criteria. SNI did not alter % nestling torn (**C**) or the overall nest score (**D**). Data represented as mean +/− SEM; n=15-21/group; unpaired t-test (Sham vs. SNI) (C) t=0.5063; p=0.6159 and (D) t=0.9642; p=0.3417.

### Chronic spared nerve injury does not alter social behaviors

Studies have shown neuropathic injury may induce robust negative affective states commonly associated with dysfunction in innate social behaviors in CD-1 mice[79]. We next sought to determine whether spared nerve injury induces deficits in natural social behaviors in C57BL/6J mice (**Figure 5A**). When exposing mice to a conspecific stranger mouse in a neutral homecage [36,67] fifteen weeks after SNI or sham surgery, there were no differences in time spent interacting (**Figure 5B**) or frequency of interactions (i.e. active attention towards stranger mouse, walking towards stranger mouse, touching, and licking) with the stranger mouse (**Figure 5C**). Similarly in a two-chamber social preference test[75,76] (**Figure 5D**) where, unlike the previously mentioned social interaction test mice, mice were given a choice between socializing or not, SNI mice spent the same amount of time on the stranger side as sham controls (**Figure 5G&H**). This indicates sixteen weeks of SNI injury does not alter valence associated with social interaction in C57BL/6J mice. As expected, there was a significant decrease within sham and SNI groups in distance travelled between the baseline and social preference test (**Figure 5E**) and no side bias prior to testing (**Figure 5F**), both of which are characteristic of this test[75,76]. Data from this experiment suggests 15-16 weeks of chronic spared nerve injury does not alter innate social behaviors themselves nor induce an aversion to social interaction.

**Figure 5:**
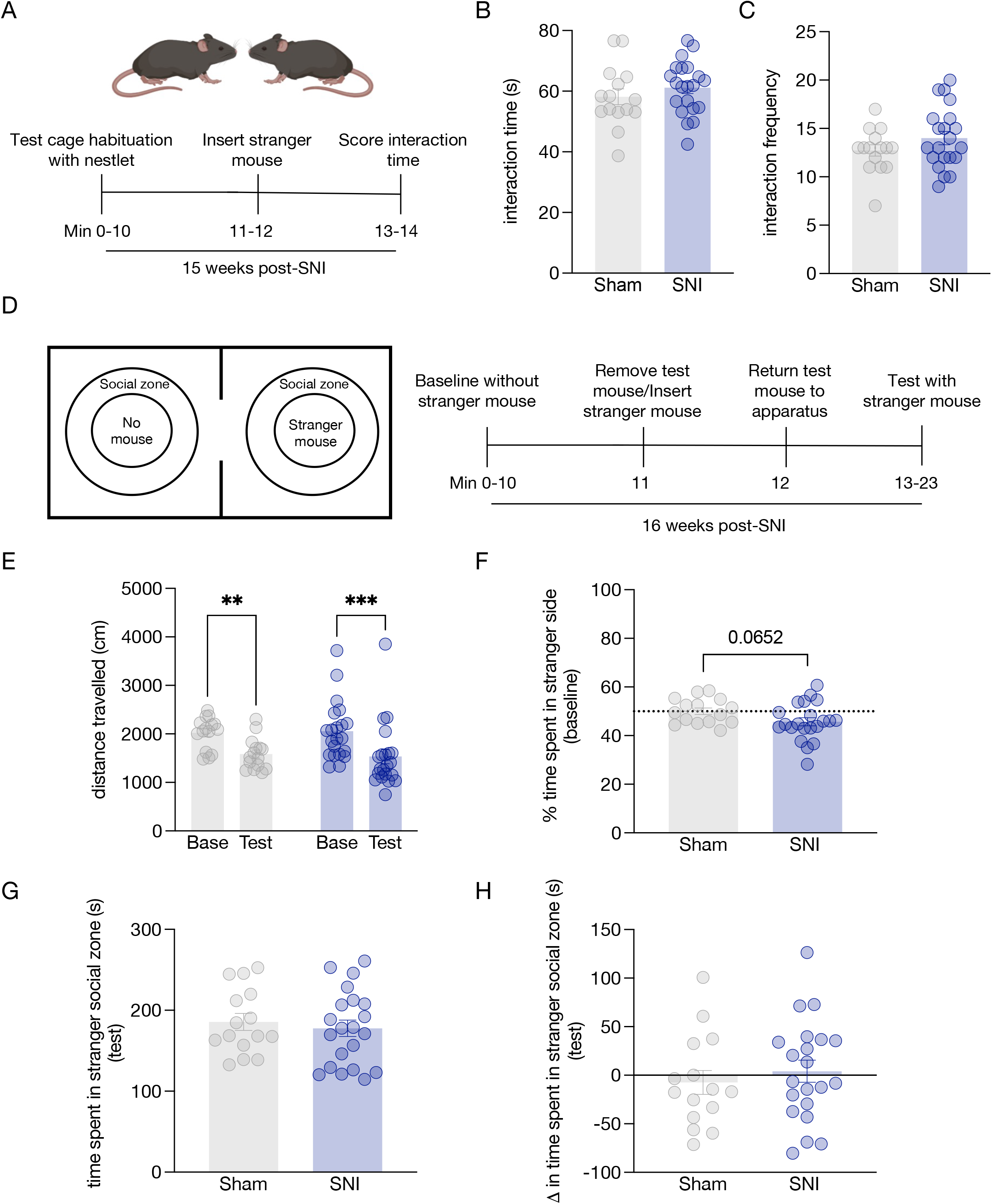
Chronic spared nerve injury does not alter social behaviors. (**A**) Experimental timeline for homecage-based social interaction. (**B**) SNI did not alter amount of interaction time between experimental mouse and stranger mouse expressed in seconds. Data represented as mean +/− SEM; n=15-21/group; unpaired t-test (Sham vs. SNI) t=0.9687; p=0.3395. (**C**) SNI did not alter frequency of interaction between the experimental mouse and stranger mouse. Data represented as mean +/− SEM; n=15-21/group; unpaired t-test (Sham vs. SNI) t=1.327; p=0.1932. (**D**) Cartoon of two-chamber social preference test and experimental timeline. (**E**) Distance travelled during the test day is decreased compared to baseline. Data represented as mean +/− SEM; n=15-21/group; two-way ANOVA with Sidak’s post hoc comparison (Sham Baseline (Base) vs. Sham test; SNI Baseline (Base) vs. SNI test) F(1,34)=0.003629; p=0.9523. SNI did not alter this reduction in distance travelled. (**F**) SNI did not significantly reduce side bias during baseline test expressed as % time spent in stranger side. Data represented as mean +/− SEM; n=15-21/group; unpaired t-test (Sham vs. SNI) t=1.906; p=0.0654. (**G**) SNI did not change the time spent in stranger side during test expressed in seconds or (**H**) the ratio of time spent in stranger side expressed as time spent in stranger side during experimental test minus time spent in stranger side during baseline test. Data represented as mean +/− SEM; n=15-21/group; unpaired t-test (Sham vs. SNI) (F) t=0.5331; p=0.5974 (G) t=0.6751; p=0.5042.

### Chronic spared nerve injury did not impede learning a fixed ratio-1 task

Preclinical assessment of pain has largely focused on sensory perception, however operant behaviors have been employed to evaluate pain-induced changes to learning and motivation [6,56,57,60,65,69]. Using open-source home-cage feeding devices (i.e., FED3) we trained mice to learn a fixed-ratio one (FR1) task. The use of operant behaviors provides critical insight into an animal’s ability to learn as well as an indicator of their motivation to complete the task. Using the FED3 devices (**Figure 6A**), animals performed the operant task overnight during the course of the 12-hour dark cycle. Most often fixed-ratio operant tasks are conducted over shorter durations and often in the light cycle. However, we hypothesized that increasing length of access to operant devices may provide more information into the overall motivational state of an animal. We previously showed that cumulative exposure to the operant task in overnight sessions can train mice on a fixed-ratio 1 task in a time efficient manner[58]. To determine whether mice successfully learned the FR1 task we set a benchmark for testing, in this case mice were deemed capable of moving to fixed-ratio 3 testing after they achieved 75% correct pokes across 3 out of the last 5 days (**Figure 6B**). Although number of days it took for each mouse to reach this benchmark criteria was variable, there was no group difference between SNI and sham animals (**Figure 6C**). To further assess learning, we compared the % correct pokes (i.e., active nosepokes as a percent of total nosepokes) between the first and final FR1 session of both sham and SNI groups. While both groups significantly increased the % of correct pokes, there was no group difference between SNI and sham animals (**Figure 6D**). In addition, we plotted the time course of pellets earned to determine whether seven weeks of neuropathic injury altered the pattern in which pellets were pursued throughout the overnight test (**Figure 6E**). There was no significant difference between sham and SNI animals in any of these learning tests. There were, however, significant group differences in the distribution of pellet retrieval time (**Figure 6F&G**) in the first and last session, suggesting seven weeks of SNI causes animals to more slowly retrieve the pellet reward, despite ultimately retrieving equivalent rewards. Finally, we quantified a proxy of meal size within the first and last test sessions and determined there were no differences in the strategy of pellet consumption between SNI and sham animal (**Figure 6H**). Taken together we can deduce that seven weeks of SNI does not alter learning or general feeding behaviors in C57BL/6J mice.

**Figure 6:**
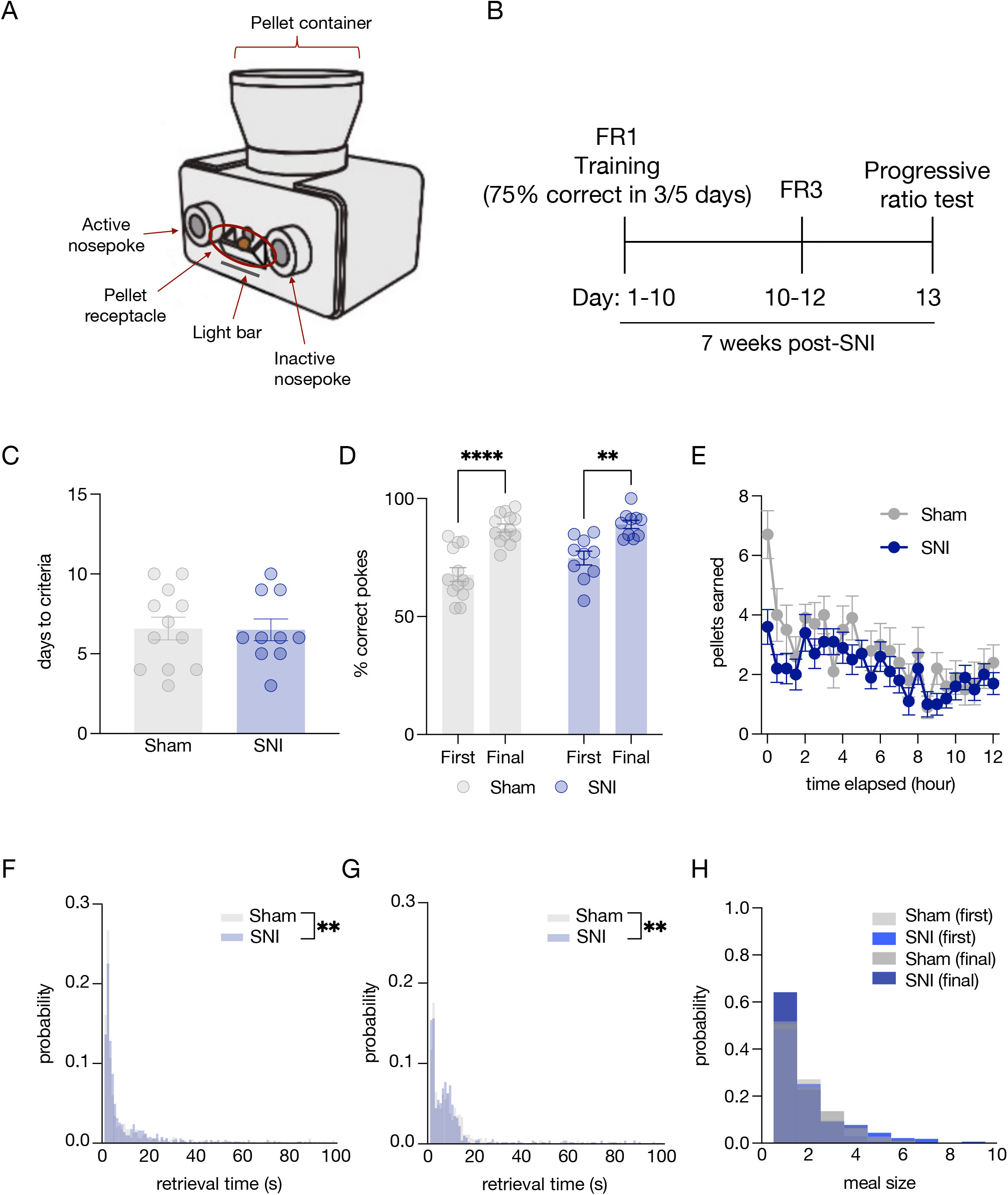
Chronic spared nerve injury does not alter performance on a homecage-based fixed ratio-1 operant task. (**A**) Cartoon identifying components of the Feeding Experimentation Device 3 (FED3). (**B**) Experimental timeline. (**C**) Neuropathic injury did not induce an inability to meet criteria in fixed ratio one task (<75% correct pokes across 3/5 days). Data represented as mean +/− SEM; n=10-12/group; unpaired t-test (Sham vs. SNI) (C) t=0.08321; p=0.9345. (**D**) Neuropathic injury did not alter learning as shown by a significant increase in % correct pokes between first and last training sessions. Data represented as mean +/− SEM; n=10-13/group; two way ANOVA with Sidak’s post hoc comparison F(1,21)=3.414, p=0.0788. € Spared nerve injury does not alter pellets earned throughout fixed-ratio one operant task. Data represented as mean +/− SEM; n=10/group; two way ANOVA with Sidak’s post hoc comparison F(1,18)=3.914, p=0.0634. Neuropathic injury also changes pellet retrieval times between sham and SNI groups in first (**F**) and last (**G**) FR1 testing session. (F) Kolmogorov-Smirnov test D=0.06161, p=0.0076. (G) Kolmogorov-Smirnov test D=0.07633, p=0.0011. Retrieval times longer than 100s were excluded from the analysis. (**H**) There was no difference in distribution of meal size between sham and SNI mice. Results of Kolmogorov-Smirnov test D=0.426, p=0.994.

### Chronic spared nerve injury alters performance during fixed ratio-3 task

Once having met the learning criteria on the FR1 task, we next sought to test whether this learning would transfer to a more difficult operant task. Mice next completed three fixed-ratio three (FR3) sessions. **Figure 7A-C** shows no difference between sham and SNI groups in distinguishing active vs. inactive pokes. This is shown as a % correct pokes. This suggests mice have learned to associate action at the active nosepoke hole to result in a sugar reward and that there is not a difference in ability to learn this association between the sham and SNI groups. Despite equivalent transfer of task learning to the FR3 task, we found a significant decrease in pellets earned in the SNI group on days one and three in consecutive FR3 sessions but not day two (**Figure 7D-F**). Additionally, SNI animals took significantly more time to earn the same number of pellets compared to sham controls (**Figure 7G**). Likewise, as in the FR1 task, neuropathic injury induced a rightward shift in pellet retrieval time distribution (**Figure 7H**), suggesting SNI mice take significantly more time to retrieve pellets upon presentation. Also, as in the FR1 task, we found no significant difference in meal sizes throughout the testing sessions between sham and SNI animals (**Figure 7I**). Together, these data suggest chronic neuropathic pain may decrease motivation for mice when more effort is required for the same reward.

**Figure 7:**
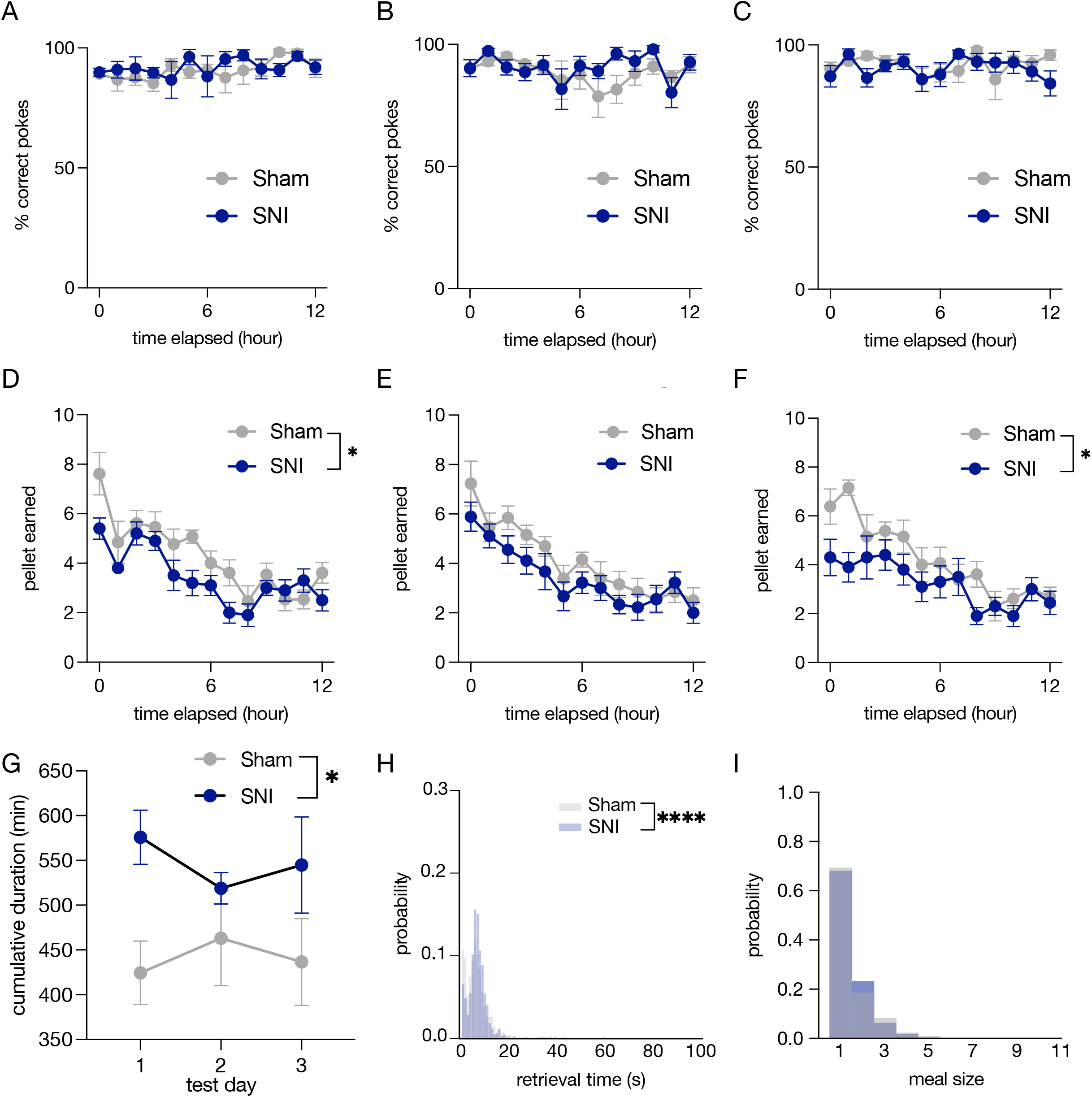
Chronic spared nerve injury reduces performance on a homecage-based fixed ratio-3 operant task. (**A-C**) We found no difference in the percentage of correct pokes during three fixed-ratio three sessions between sham and SNI groups nor a significant effect of test session. Data represented as mean +/− SEM; n=10-13/group; two-way ANOVA with Sidak’s post hoc comparison (A) F(1,21)=0.3290, p=0.5723 (B) F(1,21)=1.565, p=0.2121 (C) F(1,21)=0.6926, p=0.4146. (**D-F**) SNI decreases the number of pellets earned during three fixed ratio-3 training sessions compared to sham controls. Data represented as mean +/− SEM; n=10-13/group; twoway ANOVA with Sidak’s post hoc comparison (Sham vs. SNI) (D) F(1,21)=5.698, p=0.0265 (**E**) F(1,20)=3.888, p=0.0626 (F) F(1,21)=5.774, p=0.0256. (**G**) We found neuropathic injury increased the cumulative amount of time to reach 35 pellets across three fixed-ratio three testing sessions. Data represented as mean +/− SEM; n=10-13/group; two-way ANOVA with Sidak’s post hoc comparison F(1,20)=9.400, p=0.0264. (**H**) Relative to controls, SNI mice showed a shift in the distribution towards longer pellet retrieval times (Kolmogorov-Smirnov test D=3.802, p<0.0001; n=2247 control observations /1246 SNI observations). Retrieval latencies longer than 100s were excluded from the analysis. (**I**) There was no difference in the distribution of meal size between sham and SNI mice (Kolmogorov-Smirnov test D=0.426, p=0.994).

### Microanalysis of inter-poking behavior during fixed-ratio 3 task

After identifying potential diminished motivation in the FR3 testing shown in **Figure 7**, we became interested in determining whether strategies for completing the FR3 task diverged between sham and SNI animals. To test this question, we conducted a microanalysis of inter-poking behaviors between the three pokes (i.e., time between poke 2 and poke 1, poke 3 and poke 2, and poke 3 and poke 1) required to acquire a pellet. There was no significant difference in distribution of interpoke duration on FR3 day one (**Figure 8A,D,G**) or FR3 three (**Figure 8C,F,I**). However, there was a significant difference in inter-poke duration between all pokes on FR3 day two (**Figure 8B,E,H**). These results, together with the consistent performance across days seen in **Figure 7A-C**, suggest that sham and SNI mice likely do not have a robust difference in poking strategies to accomplish the FR3 task. This interpretation further suggests the decrease in pellets earned between sham and SNI animals shown in **Figure 7** can likely be attributed to a decrease in motivation rather than differences in poking strategy.

**Figure 8:**
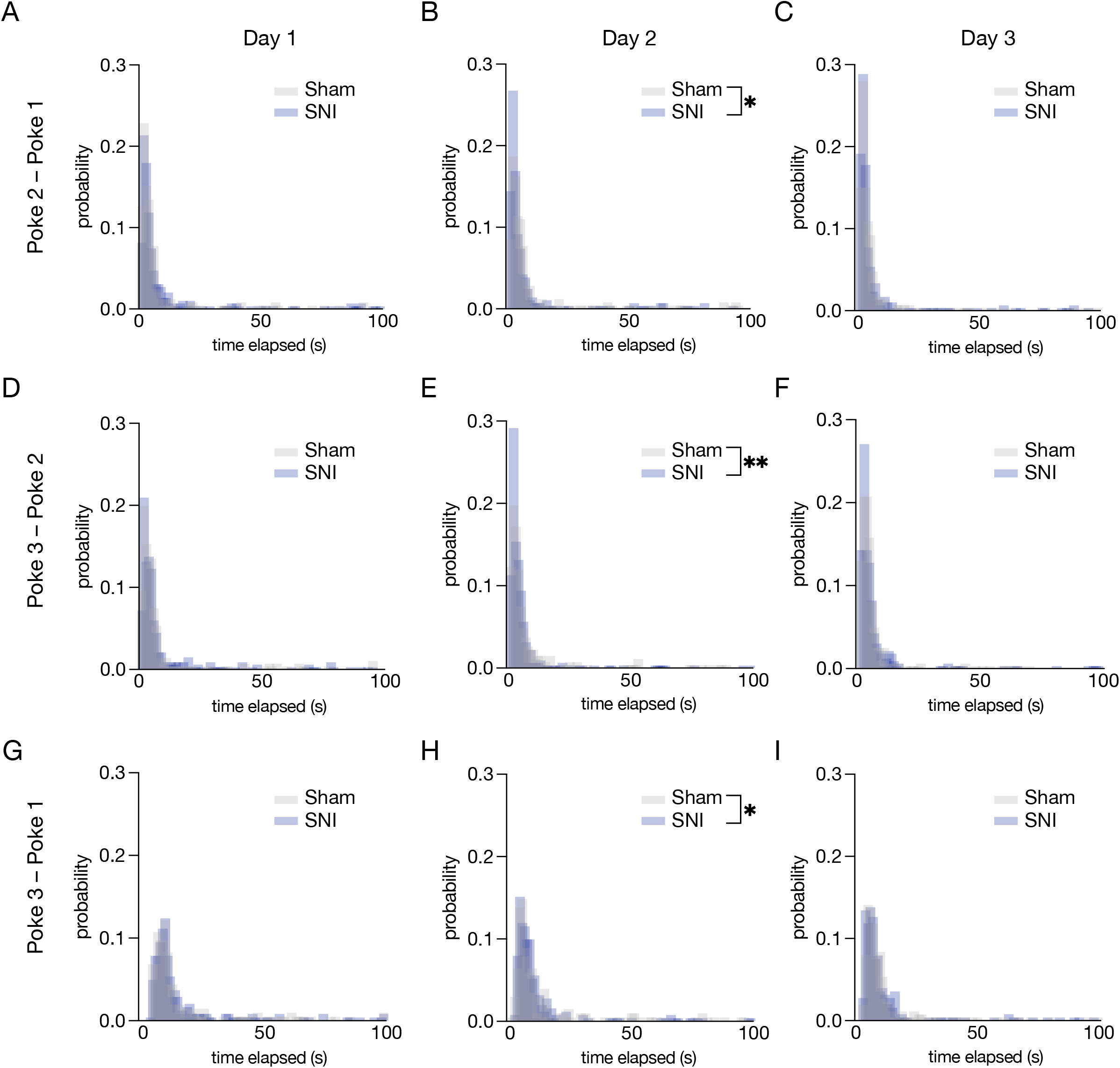
Microanalysis of poking behavior during fixed-ratio 3 task. SNI did not alter interpoke duration between pokes on day one of fixed-ratio three testing (**A, D, G**). (A) Kolmogorov-Smirnov test D=0.05892, p=0.4571, (D) Kolmogorov-Smirnov test D=0.05816, p=0.4740, (G) Kolmogorov-Smirnov test D=0.04922, p=0.6869. SNI did alter inter-poke durations on day two of fixed-ratio three testing (**B, E, H**). (B) Kolmogorov-Smirnov test D=0.1101, p=0.0174, (E) Kolmogorov-Smirnov test D=0.1166, p=0.0098, (H) Kolmogorov-Smirnov test D= 3.802, p<0.0001. Inter-poke duration was not altered in SNI mice on day three of fixed-ratio three testing (**C, F, I**). (C) Kolmogorov-Smirnov test D=0.07244, p=0.2208, (F) Kolmogorov-Smirnov test D=0.08287, p=0.1124, (I) Kolmogorov-Smirnov test D=0.09198, p=0.0576.

### Chronic spared nerve injury decreases motivation to pursue sugar rewards in a long-access progressive ratio task

To directly test whether neuropathic pain-induces motivational deficits, we next conducted an exponential progressive ratio test using the FED3 devices. In a twelve-hour overnight long-access progressive ratio task SNI mice show a significant decrease in their breakpoint (i.e., the number of pellets each animal was willing to work exponentially harder to earn) compared to sham controls (**Figure 9A**). This is further reflected by a significant decrease in leftover pokes (i.e., pokes made after the last reward that did not result in a reward) at the completion of the test (**Figure 9B**). These results suggest chronic neuropathic injury decreases motivation to perform on a long-access progressive ratio task. **Figure 9C** represents the % correct pokes across the task to confirm mice are still performing the task effectively and not exploring alternative operant strategies, there were no differences between sham and SNI groups. Additionally, as in the prior operant tasks, there was no difference in meal size between sham and SNI groups (**Figure 9D**). When examining the number of pokes each hour, SNI mice show an overall decrease in number of pokes across the entirety of progressive ratio test (**Figure 9E**). Likewise, SNI mice earn significantly fewer pellets each hour throughout the twelve-hour progressive ratio task (**Figure 9F**). **Figure 9G** shows significant differences in pellet retrieval time distribution between SNI and sham controls during the progressive ratio test, similar to what we observed in the FR1 and FR3 tasks. SNI animals also had a significant injury-induced rightward shift in inter-pellet interval distribution (**Figure 9H**). This suggests nerve injured mice dedicate less consolidated effort to retrieve a subsequent pellet. Altogether, these data indicate SNI mice have decreased performance on our long-access progressive ratio task consistent with a general decrease in motivation.

**Figure 9:**
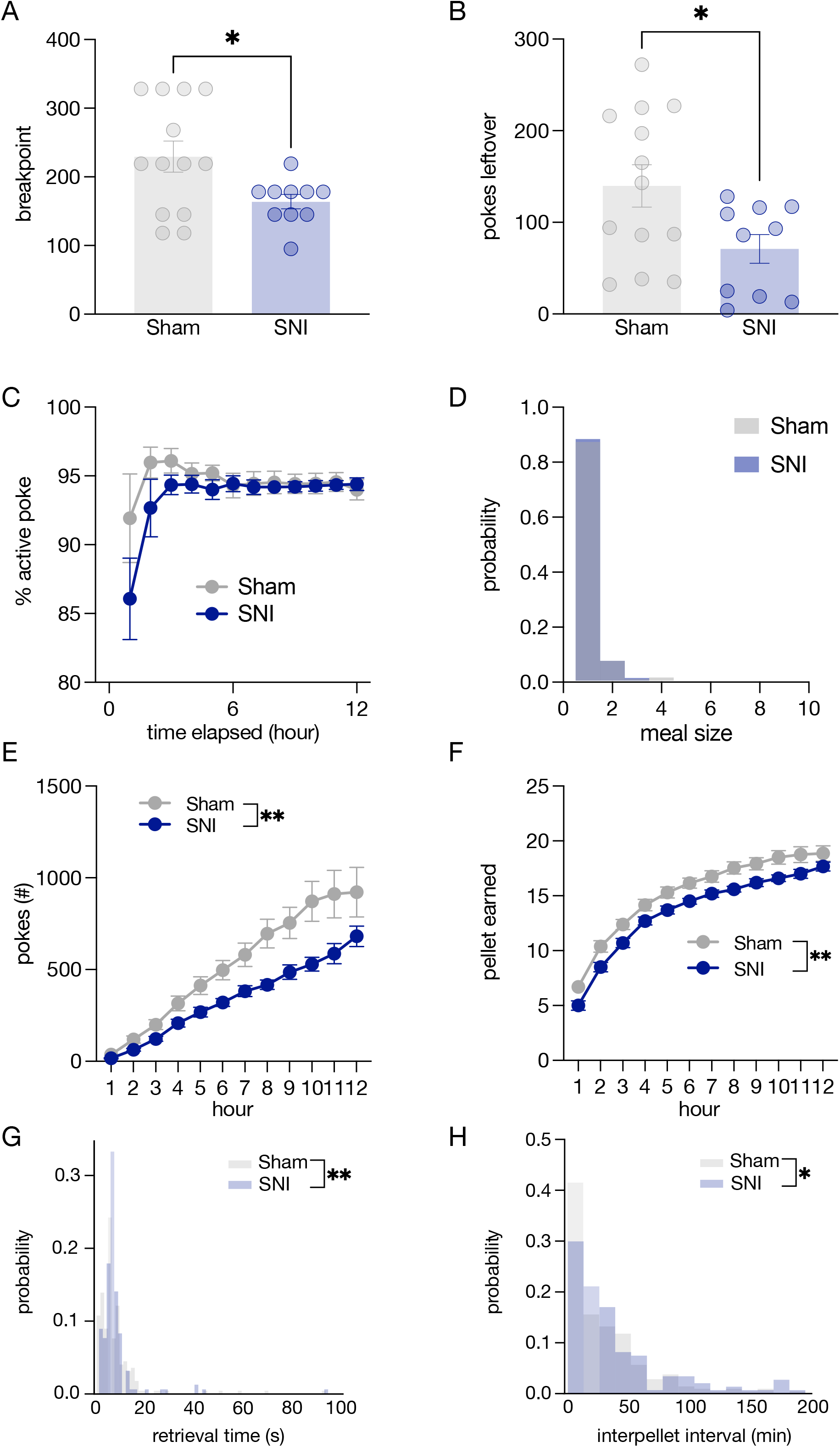
Chronic spared nerve injury decreases motivation for a sugar pellet on a homecage-based, long-access progressive ratio task. (**A**) SNI decreases breakpoint compared to sham controls. Data represented as mean +/− SEM; n=10-13/group; unpaired t-test (Sham vs. SNI) t=2.382; p=0.0268. (**B**) SNI also decreases the number of pokes following the final rewarded ratio (i.e., leftover pokes) compared to sham controls. Data represented as mean +/− SEM; n=10-13/group; unpaired t-test (Sham vs. SNI) t=2.312; p=0.0310. (**C**) There was no effect of neuropathic injury on % correct pokes throughout the progressive ratio test. Data represented as mean +/− SEM; n=10-13/group; two-way ANOVA (Sham vs. SNI) F(1,21)=0.5005, p=0.4871. (**D**) There was no difference in meal size between sham and SNI groups during progressive ratio test. Kolmogorov-Smirnov test D=0.25, p=0.998. (**E**) Neuropathic injury induced a significant decrease in the number of correct pokes completed throughout the progressive ratio test. Data represented as mean +/− SEM; n=10-13/group; two-way ANOVA (Sham vs. SNI) F(1,21)=9.338, p=0.006. (**F**) Spared nerve injured animals also showed a significant decrease in pellets earned throughout the progressive ratio test. Data represented as mean +/− SEM; n=10-13/group; two-way ANOVA (Sham vs. SNI) F(1,21)=8.263, p=0.0091. (**G**) There was no significant difference in retrieval times of earned pellet in the progressive ratio test between sham and SNI groups. Retrieval times greater than 100s were excluded from the analysis. Kolmogorov-Smirnov test D=0.1677, p=0.0072. (**H**) Spared nerve injured animals exhibit a right shifted distribution in time between pellets earned. Kolmogorov-Smirnov test D=0.1549, p=0.0198.

### Integrated behavioral scores provide emergent pain-related outcomes after SNI

The bulk of this study was designed to assess how sensitive commonly used behavioral assays for modeling features of neuropsychiatric disorders are to detecting chronic nerve injury-induced changes to negative affect. With this approach most of these commonly used assays did not have statistically significant mean differences between SNI and Sham animals (**Figures 2-5**). The exception to that was in the more challenging operant tasks (i.e., FR3 and PR; **Figures 6-9**). Prior work has demonstrated that integrating performance across various behavioral assays can lead to more sensitive and reliable measures of the overall so-called ‘emotionality’ of preclinical rodent models[9,12,35,78]. Whether this same approach would apply for injury-induced negative affect is unclear. To determine whether a normalized ‘emotionality’ behavioral z-score could reflect chronic neuropathic injury, we next tested this concept by normalizing each individual output from a single behavioral assay (e.g., % time in center and distance travelled in the open field; **Table 1**) into a single normalized assay score. Then, we integrated these separate behavioral outputs into a common, normalized behavioral z-score that encompassed each assay. Due to the limitation of some animals performing certain assays and not others, these scores were generated separately for each cohort of animals (**Table 1**). Examples of assays included in compiling emotional z-scores are shown in **Figure 10A&E**. For Cohort 1 (which included outputs from OFT, EPM, sucrose preference, tail suspension, and the operant tasks), this normalized behavioral score was significantly higher for SNI animals compared to Sham controls (**Figure 10B**). Furthermore, this behavioral z-score was significantly negatively correlated with mechanical thresholds taken the week before these tests began (**Figure 10C**). Normalized PR performance, FR3 testing, and sucrose preference test were also significantly positively correlated with the normalized behavioral z-score (**Figure 10D**). Cohort 2 (which included outputs from palatable food choice, nest building, and social tests) showed no significant difference between SNI and Sham controls (**Figure 10F**) or any significant correlations with mechanical thresholds (**Figure 10G**). Interestingly, normalized behavioral z-scores calculated from Cohort 2 were positively correlated with outputs from chocolate preference, social preference, and social interaction tests (**Figure 10H**). Though not represented graphically, a number of significant correlations also appeared between normalized behaviors including a positive correlation between EPM and sucrose preference (r = 0.546, p =0.006) and FR3 and PR (r= 0.587, p = 0.003) as well as a negative correlation between PR and TST (r=-0.460, p=0.027). Together these results suggest that integrated behavioral scores, especially those including approach-avoidance conflict, coping behaviors, and motivational tasks, may synergize to provide useful metrics for neuropathic injury-induced negative affect. Furthermore, it is worth considering that a lack of statistical significance on single behavioral output could be misleading for interpreting negative affective outcomes from neuropathic injury.

**Figure 10:**
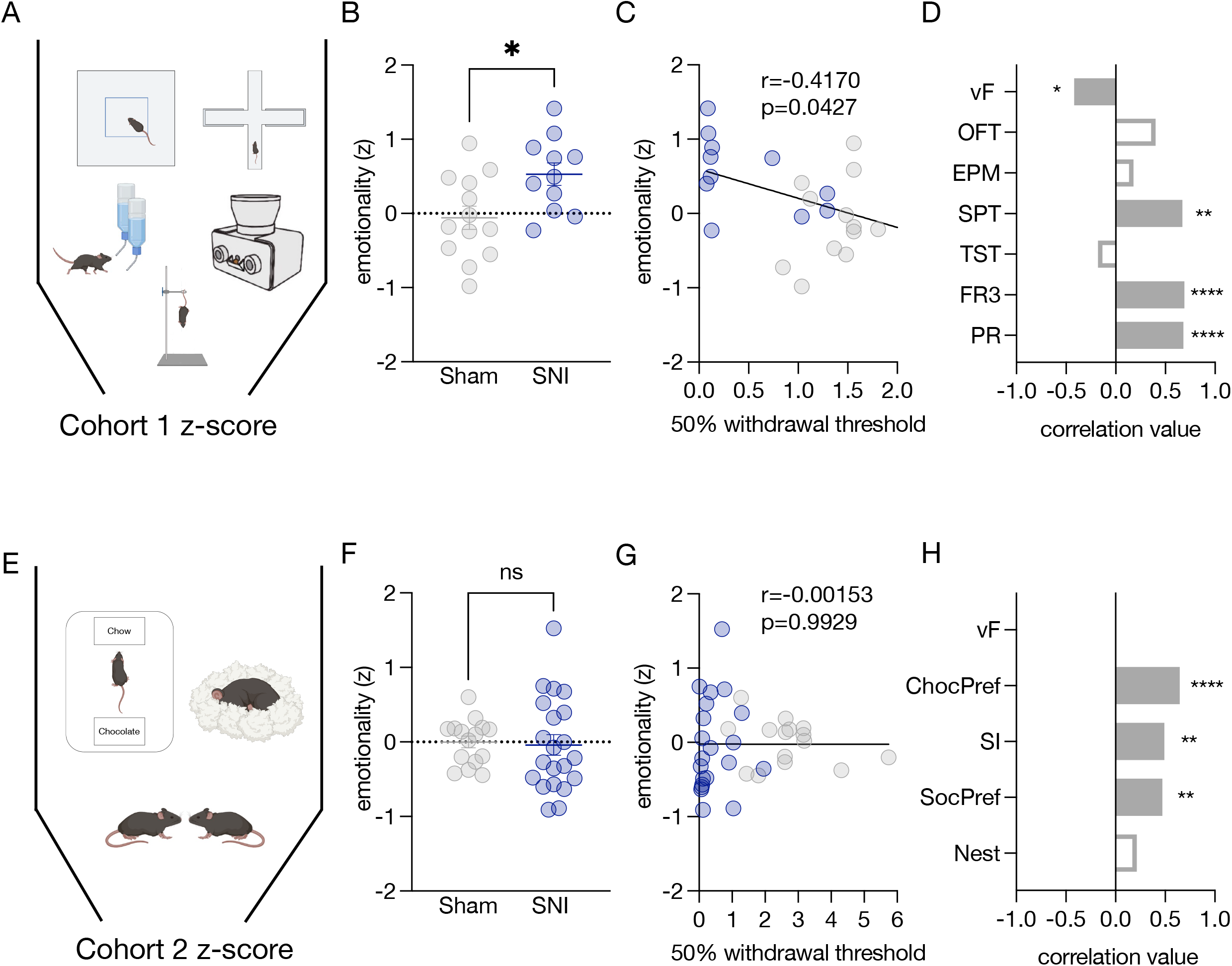
Chronic spared nerve injury induces increase in emotionality z-score associated with negative affect. (**A**) Cartoon showing behavioral assays included in compiling cohort 1 emotionality z-score. (**B**) Cohort one spared nerve injured mice exhibit overall increased emotionality z-scores. Data represented as mean +/− SEM; n=11-13/group; unpaired t-test, t=2.676, p=0.0138. (**C**) Data from cohort one shows a negative relationship between emotionality z-score and von Frey mechanical sensitivity measurements. Data shown as simple linear regression, ŕ=0.1738, p=0.0427. (**D**) Emotionality z-score is significantly positively correlated with sucrose preference test, fixed-ratio three testing, and the progressive ratio test. Pearson correlation test; sucrose preference test, r=0.674, p=0.003; fixed-ratio three, r=0.691, p=0.0002; progressive ratio test, r=0.68, p=0.0003. (**E**) Cartoon showing behavioral assays included in compiling cohort 2 emotionality z-score. (**F**) There was no significant effect of neuropathic injury on emotionality z-score in cohort 2 mice. Data represented as mean +/− SEM; n=15-21/group; unpaired t-test, t=0.2286, p=0.8206. (**G**) Emotionality z-scores did not exhibit a linear relationship to von Frey mechanical sensitivity scores. Data shown as simple linear regression, r^2^= 0.0000023, p=0.9929. (**H**) Emotionality z-score was positively correlated with social preference test, social interaction test, and chocolate preference test. Pearson correlation test; social preference test, r=0.472, p=0.004; social interaction, r=0.492, p=0.002; chocolate preference, r=0.646, p=0.00002.

## Discussion

Chronic pain remains a detrimental issue worldwide. Reliability of preclinical behavioral assays is critical to discovering the fundamental neurobiology needed to solve this complex issue. Preclinical models of pain-induced negative affect, however, have unique reproducibility and translational challenges[17,44,65,74,82]. Differential reproducibility of pain-induced negative affect between labs appears to largely arise from a variety of critical factors including behavioral assay protocols, animal subject environment prior to experimental testing, species, strain, sex, breeding strategies, and sex of the experimenter[17,24,44,52,60,77,83]. Here we aimed to clearly document pain-induced negative affect arising from one particular neuropathic injury (i.e., SNI) in one particular mouse strain (i.e., C57BL/6J) by assessing multiple behavioral outputs per animal and attempting to identify relationships between these behaviors that have not been previously tested. Pain-induced negative affect is typically thought to arise on longer timescales with the onset of chronic pain[2,44,84]. While identifying an exact timeframe for the onset of chronic pain is difficult, some attribute chronic pain onset to generalized central sensitization[46]. Following SNI, in our studies, we were able to reliably track mechanical sensitivity and reported no contralateral hypersensitivity in the earlier timepoints of spared nerve injury. However, interestingly we report mechanical allodynia on the contralateral side after 26 weeks of injury (**Figure 1D**). This suggests some degree of central sensitization at the 26-week timepoint; however, we cannot directly attribute this effect to an increase in negative affective behavior considering all behavioral testing was completed prior to acquiring this 26 week von Frey timepoint. Future studies dissecting the discrete time course of this contralateral hypersensitivity onset could prove useful in understanding the potential relationship between central sensitization and pain-induced negative affect.

As with any experiment, variability in pain-induced negative affect across labs and assays should be carefully considered. While some factors might be difficult to pinpoint, different protocols should be clearly identifiable. In our study, we showed no alterations in nesting behavior as a result of neuropathic injury (**Figure 4**). Notably however, there is a wide range of nestlet testing protocols each factoring in unique nest characteristics to configure a cumulative nest score[33,42]. Additionally, nestlet test duration and time of day is not consistent between protocols[32]. It is therefore possible, that using a different nesting protocol following SNI could reveal a clear phenotype as others have already shown across different pain-related models[3,9,43,47]. The same could be said for our sucrose preference results (**Figure 3A**). There are many different sucrose preference protocols in the literature[28,41,45,86]. Typically points of divergence in sucrose preference protocols include sucrose concentration, habituation paradigms, and counterbalancing access to water and sucrose[61]. It is worth noting that different protocols have shown significant results following injury models[86]. Furthermore, we show no effect of neuropathic injury in social interaction or social preference (**Figure 5**). Critically, however, our mice were all approximately 23 weeks of age during social testing. Previous studies have shown that age is a critical variable in the output of social behavioral assays in mice[14]. Older mice tend to spend less time interacting socially so it is conceivable that SNI could drive changes in social behaviors at earlier ages. In this study, however, we would expect any age-related effects to equally impact our sham controls.

Many groups have used traditional, ‘Skinner box’-based operant tasks to determine whether pain models affect learning and motivated responding. Depending on the type and duration of injury, the operant task tested, and the species or strain used, these models have produced mixed results[6,29,56,57,60,64,65,69]. Here we report that, in male and female C57BL/6J mice, SNI reduces operant responding for sugar pellets in a novel overnight, homecage fixed-ratio 3 and progressive ratio tasks. SNI also slows sugar pellet retrieval in all the operant tasks we tested compared to sham controls. Given that overall locomotion was not decreased in exploratory behaviors (**Figure 2C&F**), this increased latency in pellet retrieval could be related to overall motivation to complete each aspect of the task.

For these operant tasks, mice were tested in a dedicated homecage outfitted with FED3 devices in 12-hour, overnight sessions. These open-source devices are easy to build and inexpensive to acquire[58], meaning that studying pain-induced decreases in motivated behavior is now accessible across the funding landscapes of different laboratories. A potential limitation of the training procedure we used, however, is the variable length of time mice take to learn a task, which can interact with habituation to the novel test environment and handling required to move mice in and out of the chambers. This approach minimizes these concerns by giving long-access to the operant device (i.e., FED3) for time-efficient learning (**Figure 6C**). Furthermore, the point in the circadian cycle of testing and level of food consumption immediately prior to the task will also impact motivation in food-rewarded tasks[8,40,58]. To mitigate these concerns, mice were tested on overnight sessions, starting at the beginning of the dark cycle; session times were kept consistent between and across animals and no food restriction was used. While this approach clearly and efficiently identifies changes to motivation following chronic SNI, it is worth considering expanding the scope of these studies. Namely, future studies that employ ‘closed-economy’ behavioral paradigms, in which mice have access to the device 24 hours a day, 7 days a week as their only source of food, will allow for more explicit control of circadian and satiety variables, and will allow for more complex behavioral tasks to assess cognitive changes in SNI mice. Moreover, closed-economy paradigms would allow for longitudinal measurements of behavior from baseline through the induction and stabilization of the chronic pain state following SNI with minimal experimenter intervention[7,30,48,67]. These longitudinal experiments would shed light on whether variability in performance emerges because of SNI, or if individual differences in performance exist prior to the injury and are exacerbated with the induction of chronic neuropathic pain.

In addition to progressive ratio testing, other operant tasks have been used to identify cognitive impairment and attentional deficits as a result of chronic neuropathic injury[21,38]. In particular, one important prior report showed that spinal nerve ligated rats produce behavioral bursts of lever-pressing in a modified operant task with a choice for either one reward or four rewards. In our study, we quantified inter-poke interval variability in our FR3 operant task to determine whether SNI drives similar differences in poking strategies in mice (**Figure 8**). While we did see increased probability of shorter inter-poke durations in SNI mice during FR3 testing this effect only occurred in two out of three test days. This day-to-day variability suggests that this measure may not be a robust affective outcome following SNI in mice. It would be useful for future studies to identify whether the same bursting pattern observed in rats is consistent in mice in the same operant choice task.

Importantly, rodents used in this study were all male and female C57BL/6J mice. This point is critical to the overall interpretation of the experiments because sex and strain differences in neuropathic pain-induced negative affect are well known across species in the literature[24,60,82]. For example, one study found Balb/c mice increased exploration of the center zone in the OFT, a proxy for decreased anxiety-like behavior, one month of spared nerve injury. On the contrary, C57BL/6J mice had no effect of neuropathic injury at the same timepoint[82] – similar to the results we show here in **Figure 2A-C**. Another important factor to consider when evaluating prior studies is the origin of the animal subjects. In Urban et al, 2011[82] both the Balb/c and C57BL/6J mice were purchased and transferred to the experimental facility, whereas mice in our study were bred locally and transferred to our experimental facility. In each of these cases there was some ground transportation, which is a well-established stressor for mice[73,80], and could set these studies apart from groups that are able to breed and test animals in the same facility. Counter to this point, however, others have shown that long-term SNI did cause anxiety-like behavior in mice that had been purchased and transferred[74,86], but these C57BL/6 were from different vendors and thus may have substantial genetic differences between these populations[16]. Regardless, it is undoubtedly critical for investigators to clearly track and report the origin and stress history of animals in these types of studies.

Finally, we aggregated our data to report normalized emotionality z-scores[35]. These scores reflect a generalized affective state rather than declaring an affective state based on a single behavioral assay. These emotionality z-scores reflect a consolidation of several outputs from several behavioral assays. Despite our studies presenting with largely ‘negative data’ that do not, alone, imply neuropathic injury-induced negative affect on an individual assay basis, these normalized scores suggest we were still able to identify a negative affective state following SNI (**Figure 10B**). It is important to note these emotionality z-score outputs do not seem to be driven exclusively by behavioral assays that were individually statistically significant, suggesting there is likely information to gain from ‘negative data’ (i.e., not statistically significant) outputs on individual behavioral assays. Altogether, we used a battery of behavior tests often used in the study of negative affective behavior to determine the neuropsychiatric impact of long-term neuropathic injury in C57BL/6J mice. While our most notable single results were from homecage-based long-access operant tasks, we have also demonstrated that normalized behavioral scores across a battery of assays were able to identify a generalized negative affective state induced by chronic spared nerve injury. As a result, we encourage others to consider using similar approaches when studying pain-induced negative affect.

## Acknowledgements

We thank the other members of the Al-Hasani and McCall labs for helpful feedback on this project. This work was supported by the National Institutes of Health (R01NS117899, J.G.M.; F31NS124301, M.R.N.; T32DA007261, J.B.; R01NS123070, R.A.; R01DA049924, M.C.C.), the Hetzler Foundation (Pilot grant to M.C.C.), and the Rita Allen Foundation (M.C.C. and J.G.M.) with help from the Open Philanthropy Project (J.G.M.). We would like to acknowledge biorender.com for figure cartoons.

## Author contributions

M.R.N and J.G.M conceived the project and designed the detailed experimental protocols. M.R.N. and J.B. performed the mouse experiments. M.R.N., L.J.B., J.B., G.B., Y-.H.C, S.S.D, M.K.M., M.M.C., and J.G.M performed the investigation and analyzed the data. M.R.N., L.J.B., J.B., R.A., M.C.C., and J.G.M wrote the paper. M.R.N., L.J.B., J.B., Y-.H.C., M.K.M., R.A., M.C.C., and J.G.M edited the paper. M.R.N. and J.G.M acquired funding. R.A., M.C.C., and J.G.M. provided research supervision. J.G.M. led overall project administration. All authors discussed the results and contributed to revision of the manuscript.

## Conflict of Interest

The authors declare no conflicts of interest.

## Notes

### Competing Interest Statement

The authors have declared no competing interest.

